# The mitochondrial *orf117Sha* gene desynchronizes pollen development and causes pollen abortion in the Arabidopsis Sha CMS

**DOI:** 10.1101/2024.01.17.575984

**Authors:** Noémie Dehaene, Clément Boussardon, Philippe Andrey, Delphine Charif, Dennis Brandt, Clémence Gilouppe Taillefer, Thomas Nietzel, Anthony Ricou, Matthieu Simon, Joseph Tran, Daniel Vezon, Christine Camilleri, Shin-ichi Arimura, Markus Schwarzländer, Françoise Budar

## Abstract

Cytoplasmic male sterility (CMS) is of major agronomical relevance in hybrid breeding. In gametophytic CMS, abortion of pollen is determined by the grain genotype, while in sporophytic CMS, it is determined by the mother plant genotype. While several CMS mechanisms have been dissected at the molecular level, gametophytic CMS has not been straightforwardly accessible. We used the gametophytic Sha-CMS in Arabidopsis to characterize the cause and process of pollen abortion by implementing *in vivo* biosensing in single pollen and mitoTALEN mutagenesis. We obtained conclusive evidence that *orf117Sha* is the CMS-causing gene, despite distinct characteristics from other CMS-genes. We measured the *in vivo* cytosolic ATP content in single pollen, followed pollen development and analyzed pollen mitochondrial volume in two genotypes that differed only by the presence of the *orf117Sha* locus. Our results show that the Sha-CMS is not triggered by ATP deficiency. Instead, we observed desynchronization of a pollen developmental program. Pollen death occurred independently in pollen grains at diverse stages and was preceded by mitochondrial swelling. We conclude that pollen death is grain-autonomous in Sha-CMS and propose that mitochondrial permeability transition, which was previously described as a hallmark of developmental and environmental-triggered cell death programs, precedes pollen death in Sha-CMS.

**Highlight:** The Arabidopsis CMS-causing gene *orf117Sha* does not limit pollen ATP supply. Pollen-centered approaches show desynchronization of development and mitochondrial swelling before pollen death, which occurred at diverse stages.

## Introduction

Cytoplasmic male sterility (CMS), a source of reproductive polymorphism in Angiosperms, is genetically determined by both nuclear and maternally inherited mitochondrial factors (Chittenden and Pellew, 1927; Dufaÿ *et al*., 2014). Mitochondria contain a CMS-causing gene, which prevents the plant from producing functional pollen, unless its nuclear genome possesses restorer-of-fertility (Rf) gene(s). In nature, the presence of a mitochondrial CMS gene in a population where Rf is absent or in segregation with non-restorer allele(s) results in female (male sterile) individuals among hermaphrodite congeners, a coexistence named gynodioecy (Darwin, 1877). When Rf genes are fixed, the CMS becomes cryptic: male sterility is no longer phenotypically expressed, but can be observed again after introduction of the cytoplasm into a non-restorer nuclear background.

CMS first attracted the interest of plant breeders in the 1940s, to facilitate the field-scale production of hybrid seeds and exploit heterosis in elite varieties (Jones and Clarke, 1943). In the vast majority of cases, CMSs used in agriculture rely on naturally occurring mitochondrial CMS genes, most often cryptic in the species, subspecies, or cultivar of origin. Studies to identify CMS-causing genes and understand the mechanism(s) of pollen abortion have been conducted mainly in cultivated plants, but also in Mimulus (Case and Willis, 2008), Silene (Stone *et al*., 2017), and wild beet (Meyer *et al*., 2018). Mitochondrial CMS genes have been identified in more than 30 CMSs in approximately 20 species (reviewed by Chen & Liu (2014), Toriyama (2021), Xu *et al*. (2022)). Several features common to many, but not all, identified CMS-genes were drawn from the comparison of these genes (Hanson and Bentolila, 2004; Horn *et al*., 2014; Chen *et al*., 2017; Kitazaki *et al*., 2023), and sometimes used to drive the search for mitochondrial CMS-causing genes: a chimeric sequence comprising identifiable parts of conserved mitochondrial genes and sequences of unknown origin; association and co-transcription with a conserved mitochondrial gene; expression profile modified by the action of nuclear Rf, most often at the RNA level; a predicted protein product containing hydrophobic transmembrane domain(s). While mitochondrial-CMS genes were first thought to be unique to a species or population, some were found to have evolved variants or related mitochondrial CMS genes within plant taxa, mainly in the Brassicaceae family (L’Homme *et al*., 1997; Yamagishi and Terachi, 2001; Yamagishi *et al*., 2021b,*a*) and the rice genus (He *et al*., 2020; Zhang *et al*., 2022).

Two genetic types of CMS and restoration have been distinguished since the early genetic studies of CMS (Bucher, 1961; Duvick, 1965; Laughnan and Gabay-Laughnan, 1983). In gametophytic CMS, pollen abortion is determined by the pollen grain genotype; in an individual heterozygous for the Rf, pollen grains that carry the restorer Rf allele survive, while those carrying the non-restorer allele die. In contrast, in sporophytic CMS, pollen abortion is determined by the mother plant genotype because pollen abortion results from a default in the diploid maternal tissue of the anther, most often the tapetal cell layer. Sporophytic Rf genes are usually dominant, and heterozygotes produce 100% viable pollen. It seems reasonable to assume that tapetum and pollen abortions are triggered by different mechanisms in sporophytic and gametophytic CMSs. Several studies support this hypothesis. In the sporophytic *Brassica napus nap*-CMS, *orf222* expression blocks microsporangium development at an early and specific stage, but in anther sites where sporogenesis does take place, once this stage is passed, *orf222* expression does not impair male gametogenesis or the production of functional pollen (Geddy *et al*., 2005). In rice, the genetic feature (*i.e.* sporophytic or gametophytic) of each CMS was conserved when the 35SCaMV promoter, expressed in many sporophytic organs but also in pollen in this species, was successfully used to drive the expression of the sporophytic FA-CMS (Jiang *et al*., 2022) or the gametophytic BT-CMS (Wang *et al*., 2006) genes in transgenic plants.

Despite the number of identified CMS-genes, the mechanism by which the CMS-gene product triggers pollen death has only been partially elucidated in a few cases, mainly in sporophytic CMSs. Several causes were reported or proposed to trigger tapetum premature death: the CMS-protein forming a pore in the mitochondrial membrane, as in the T-CMS of maize (Levings, 1993) or the Ogura CMS in rapeseed (Duroc *et al*., 2009); interaction of the WA-CMS protein with a subunit of mitochondrial complex IV provoking a burst of reactive oxygen species (ROS) in rice; impairment of sufficient ATP production by disturbing the activity of ATP synthase, as in PET1-CMS in sunflower (Sabar et al., 2003), C-CMS of maize (Yang et al., 2022) and pepper CMS (Ji *et al*., 2013; Li *et al*., 2013). In contrast, in C5- CMS cabbage, impaired ATP synthase activity and lower ATP content were associated with delayed tapetum degeneration, also resulting in male sterility (Zhong *et al*., 2022). However, it is still unclear to what extent the different causes listed above trigger different physiological mechanisms to eventually provoke pollen death and, in many cases, how these mechanisms are restricted to the tapetal cells.

To our knowledge, the molecular mechanism of pollen abortion has been unveiled only in one CMS of the gametophytic type, the HL-CMS of rice. In this case, ORFH79, the protein product of the mitochondrial CMS-causing gene, was shown to interact with the P61 protein, a subunit of cytochrome c reductase (complex III). ORFH79-P61 interaction would explain the decreased activity of the complex, accompanied by a decrease in ATP content and an increase in ROS content, observed in mitochondrial extracts from the sterile genotype compared to the fertile one (Wang *et al*., 2013). The authors proposed that limitations in ATP supply and ROS accumulation trigger retrograde signals that stop pollen development at the G1/S stage of bicellular pollen in HL-CMS plants (Wang *et al*., 2013).

In the model species *Arabidopsis thaliana*, we discovered a cryptic gametophytic CMS by crossing distant natural variants (Gobron *et al*., 2013; Simon *et al*., 2016). All male-sterility inducing cytoplasms were closely related to that of Shahdara (Sha), from Tajikistan. In contrast, the cytoplasm of Kz-9, from Kazakhstan, which is closely related to that of Sha, did not induce male sterility (Gobron *et al*., 2013; Simon *et al*., 2016). Based on restriction length polymorphisms and DNA hybridization comparing Sha and Kz-9 mitochondrial genomes, we identified *orf117Sha*, a novel mitochondrial gene unique to CMS-inducing cytoplasms, as a candidate for the CMS-causing gene. *orf117Sha* is not associated with any essential mitochondrial gene but its 5’ untranslated region is identical to that of the *cob* gene (Gobron *et al*., 2013; Durand *et al*., 2021). The ORF117SHA protein contains no identifiable fragments of any known protein necessary for the assembly or function of mitochondrial respiratory complexes, ATP synthase, or any hydrophobic putative transmembrane region. Unexpectedly, the restoration of full fertility, either by the presence of Sha Rf genes (*i.e.,* in the Sha genotype) or by inactivation of a gene necessary for sterility, was not accompanied by a modification in the expression profile of *orf117Sha* (Durand *et al*., 2021). Nevertheless, *orf117Sha* encodes a protein 56 % identical to that encoded by *orf108*, a mitochondrial gene from *Moricandia arvensis* that induces gametophytic CMS in Brassica species and is co-transcribed with *atpA* (Ashutosh *et al*., 2008; Kumar *et al*., 2012). The Sha cytoplasm-induced CMS (Sha-CMS) is phenotypically expressed when the cytoplasm of Sha is associated with the nuclear genome of Cvi-0, from the Cape Verde Islands (Roux *et al*., 2016; Durand *et al*., 2021), a genotype noted [Sha]Cvi.

In the present study, we aimed at understanding the cause and process of pollen abortion in the Sha-CMS implementing up-to-date pollen centered approaches. We definitely demonstrate that *orf117Sha* is the causative gene for Sha-CMS. We rule out that Sha-CMS is triggered by an insufficient ATP supply by experimentally measuring the *in vivo* ATP content in developing pollen. We show that pollen development is desynchronized and that death occurs independently in pollen grains, at diverse pollen developmental stages. In addition, we report that pollen death is preceded by the swelling of mitochondria reminiscent of the mitochondrial morphology transition observed in stress-induced cell death and other cases of male sterility.

## Material and Methods

### Plant material and growth conditions

#### Plant material

All the genotypes used in this study (Table 1) are available on demand at the Versailles Arabidopsis Stock Center (https://publiclines.versailles.inrae.fr/) which provided the natural variants for this work. We use the following nomenclature: a plant carrying the cytoplasmic genomes from parent A and the nuclear genome from parent B is designated [A]B.

**Table 1:** Genotypes used in this study.

| name | Cytoplasmic genomes | Nuclear background | Fertility phenotype | reference |
| --- | --- | --- | --- | --- |
| Shahdara (Sha) | Sha | Sha | fertile | natural variant |
| Kz-9 | Kz-9 | Kz-9 | fertile | natural variant |
| Cvi-0 | Cvi-0 | Cvi-0 | fertile | natural variant |
| [Sha]Cvi | Shahdara | Cvi-0 | sterile | Roux *et al.*, 2016 |
| [Kz9]Cvi | Kz-9 | Cvi-0 | fertile | this work |
| [Sha]CviL3<sup>s</sup> | Shahdara | Cvi-0<sup>a</sup> | poorly fertile<sup>b</sup> | this work |

<sup>a</sup>See material and methods for the [Sha]CviL3<sup>s</sup> genotype.
<sup>b</sup>See Fig. 3.

The [Kz9]Cvi genotype was produced by recurrent paternal backcrosses of a F1 hybrid Kz-9 x Cvi-0. A plant from the sixth backcross was sequenced on a HiSeq3000 at the Genotoul Platform in Toulouse (France) and no residual zone of heterozygosity was detected. The [Sha]CviL3^S^ possesses the Sha cytoplasm and the Cvi-0 nuclear genome except at the bottom of chromosome 3, from 21.8 Mb to telomere, where it possesses the Sha alleles.

#### Growth conditions

Plants were routinely grown in a greenhouse in long-day conditions (16 h day, 8 h night) supplemented with artificial light (105 µE/m^2^/s) when necessary. Before direct sowing on soil, seeds were stratified in the dark at 4°C for three days in water with 0.1 % (w/v) agar. *In vitro* plantlets were grown in a culture chamber under long day conditions (16 h day, 8 h night) at 20°C. Surface sterilized seeds were sown on Arabidopsis medium (3.1 g/L Gamborg B5 medium with vitamins (DUCHEFA Biochemie), 0.08% (w/v) bromocresol purple, 1 % sucrose (w/v), 0.7 % (w/v) agar, pH 6) supplemented, if needed, with the appropriate chemical for selection. When needed, plantlets grown *in vitro* were transferred to soil after two to three weeks and cultivated under the greenhouse conditions indicated above for further analyses.

### *De novo* organelle genome sequencing, assembly and annotation

#### Preparation of samples

Organelle-enriched pellets were obtained from floral buds (50-100g) following the procedure described by Scotti et al (2001), frozen and kept at −80°C until DNA extraction.

For Illumina sequencing, DNA was extracted from the organelle-enriched pellets with the NucleoSpin Plant II kit (Macherey Nagel) following manufacturer’s instructions. For PacBio sequencing, DNA was extracted according to Mayjonade et al’s (2016) procedure.

#### Assembly of chloroplast and mitochondrial genomes

Illumina sequences were produced at The Genome Analysis Centre (UK). Libraries were sequenced with 2 × 100 bp paired-end reads. As the coverage of the organelle genomes was very high (>2000x) (Table S1), Illumina reads were sampled to keep one fourth of the reads. As reads were produced from organelle-enriched DNA samples which contain nuclear, mitochondrial and chloroplast DNA we used MetaVelvet (version 1.2; Namiki *et al*., 2012), a robust assembly tool tailored for metagenomic datasets. Two sets of parameters were required (i) the expected genomes coverage (sequencing depth) and (ii) the insert-size length. We followed the guidelines provided at <http://metavelvet.dna.bio.keio.ac.jp/MV.html#advanced> to estimate the first one (Table S1). The second one (insert size) was obtained by mapping reads to the reference TAIR 10 *Arabidopsis thaliana* genomes using bwa (Li and Durbin, 2009) (Table S1). Before assembly, nuclear sequences were filtered out: khmer was used to remove reads with low abundance k-mers (Table S2). The shell mitology pipeline (https://github.com/jos4uke/mitology-pipeline.git) was implemented to automatically perform these steps for filtering and assembling short reads sequencing samples. Tables S2 and S3 give parameters used and results obtained with the mitology pipeline, respectively. As insert sizes of Illumina libraries were short compared to the size of genomic repeats, this first assembly step produced several contigs for each genome (Table S3). We then aligned raw reads back onto the contigs to check for assembly inconsistencies. In order to scaffold contigs, the R package contigLink (https://github.com/jos4uke/contigLink) was implemented to graphically explore these alignments: inspect contig inconsistencies and exploit cross contig spanning paired-end reads information. This strategy allowed full assembly of Sha and Kz-9 chloroplast genomes, while mitochondrial genomes remained as unassembled scaffolds. In order to achieve mitochondrial genome assemblies, we used PacBio long reads.

PacBio sequencing was carried out on a single SMRT cell utilizing PACBIO RSII sequencing technology at the GeT-PlaGe sequencing platform (Toulouse). Library was constructed using SMRTbell Template Prep Kit 3.0 and adapters were added using DNA/Polymerase Binding Kit P6 v2. CANU (v1.5; Koren *et al*., 2017) was used to assemble Sha and Kz9 organelle genomes (genomeSize=500k, minReadLength=500, corOutCoverage=999). Assemblies were corrected using pilon (v1.22; Walker *et al*., 2014) using Illumina reads. The resulting contigs did not correspond to fully assembled mitochondrial genome sequences. In addition, the total length of mitochondrial contigs for Sha was small (212 kb, compared to 350 kb for Kz-9, Table S4) and several essential mitochondrial genes were not detected in Sha contigs by BLAST search, while there were present in Kz-9 ones. Nevertheless, the CANU contigs were used to drive the final assembly of Metavelvet scaffolds. After this final assembly step, Sha and Kz-9 mitochondrial genome sequences were validated in two ways. Firstly, Illumina reads were aligned on the final mitochondrial genomes and unexpected pair orientation or fragment size were systematically examined using the Integrative Genomics Viewer tool (IGV_ 2.12.3) (Thorvaldsdottir *et al*., 2013). All were compatible with recombination events between copies of repeated sequences > 350 pb present in the genomes (Table S5). IGV visualization of alignments also revealed a few small regions that were not covered by Illumina reads, but present in PacBio reads. All these regions were checked by direct sequencing of specifically designed PCR products (see Table S6 for primers). Secondly, we checked that the genome sequences were consistent with the RFLP results previously produced to compare the two mitochondrial genomes (Gobron *et al*., 2013) by predicting in silico the RFLP results with InsilicoRFLP (available at https://forgemia.inra.fr/ijpb-bioinfo/public/insilicorflp.git).

#### Annotation

Chloroplast genome annotation was realized using GeSeq (Tillich *et al*., 2017, last accessed 15^th^ May 2023); the resulting annotated genomes were visualized with OGDRAW (Greiner *et al*., 2019, last accessed 21^th^ July 2023). Mitochondrial genome annotation was realized using GeSeq, last accessed 28^th^ February 2023. Annotation of trans-spliced genes was manually corrected. In addition, we used the Getorf tool of the EMBOSS program (version 6.6.0) to predict open reading frames (orfs) potentially encoding peptides at least 100 amino-acid long in Sha, Kz-9 and Col-0 (accession number: NC_037304). After discarding orfs corresponding to already annotated genes with bedtools (version 2-2.29.0), the remaining orfs were added to the annotation of Sha and Kz-9 mitochondrial genomes. We also annotated repeated sequences above 100 base pairs (Table S5) using YASS tool on line (https://bioinfo.univ-lille.fr/yass/index.php; Noé and Kucherov, 2005) for aligning each genome against itself. The resulting annotated genomes were visualized with OGDRAW, last accessed 21^th^ July 2023.

### Production and analyses of transgenic plants with nuclear expression of *orf117Sha*

In order to produce ORF117SHA and target it to the mitochondrial matrix of fertile plants, the *orf117Sha* coding sequence was fused to the mitochondrial targeting sequence from *Nicotiana plumbaginifolia* β-ATPase (Logan and Leaver, 2000), amplified from the mt-roGFP1 construct (Schwarzlander *et al*., 2008) (see Table S6 for primers) and introduced into the pUB-DEST Gateway vector (Grefen *et al*., 2010). In this construct the gene was under the control of the *UBQ10* (*AT4G05320*) promoter, which is active in developing pollen from the bicellular stage (validated using YFP fluorescence from the cATeam construct, Fig. S1). The construct was introduced into Arabidopsis Cvi-0 plants by floral dipping. Transformants were selected on sand irrigated with water containing Basta herbicide (7.5 mg.L^-1^ phosphinotricin). T1 plants were transferred to soil, verified by genotyping and grown until flowering.

Pollen viability of T1 plants was examined after Alexander staining (Alexander, 1969) of mature pollen just before anther dehiscence. The percentage of viable pollen was visually inspected and scored by allocating marks between 0 (all pollen dead) and 4 (all pollen viable). The mark for each plant was the average of the anthers from two flowers. Seeds that developed after selfing were harvested from T1 plants and transmission of the transgene was analyzed in the T2 generation either by *in vitro* screening of phosphinotricin (10 mg.L^-1^) resistant plants or by PCR. Between 90 and 300 progenies were screened for each T2 family and a Chi square test was applied to detect a bias in the transmission of the transgene.

### Production and analyses of mitochondrial mutants by mitoTALEN

#### Generation of mitochondrial mutants

Target sequences for mitoTALENs in the *orf117Sha* gene were selected by using TAL effector nucleotide targeter 2 old version (https://tale-nt.cac.cornell.edu/node/add/talen-old). The 15bp apart sequences GCCACCCCGTTGGACC, on plus strand, and CCCCGAGGATATCTTG, on minus strand, were set as targets for the left and right of TALEN pair mTAL1, both of which were checked as unique sequences in the mitochondrial genome of Sha. The sequences TACAAAACAACGCTAT, on plus strand, and TGCAGTTTCATACACT, on minus strand, were set for mTAL2 (Fig. S2). The platinum TALEN ORFs designed to recognize the target sequences were assembled by platinum gate assembly kit (Addgene) (Sakuma *et al*., 2013), and transferred into Ti plasmids by multisite Gateway LR reaction (Thermofisher) for simultaneous expression of right and left platinum TALENs fused to the Arabidopsis delta prime subunit mitochondrial presequence, under the control of the promoter of Arabidopsis *RPS5A* gene, as described previously (Kazama *et al*., 2019; Arimura *et al*., 2020). All plasmids required for assembling the tandem expression vectors for mitoTALENs in Ti plasmids are available from Addgene (Arimura, 2021). The Ti plasmid backbone is originally from a VIB Gateway vector, pK7WG2 (Karimi *et al*., 2002), which was modified to add the presequence of the Arabidopsis mitochondrial ATP synthase delta prime subunit and the Oleosin-GFP construct for the selection of T1 seeds (Shimada *et al*., 2010). Each construct carrying one pair of mitoTALENs targeting *orf117Sha* was introduced into [Sha]CviL3^S^ plants (Table 1) by floral dipping. Transformed T1 seeds were selected on the basis of GFP fluorescence. Ten seeds that showed strong GFP signal were sown on soil in the greenhouse, and the presence of the construct was verified by PCR on seedling total DNA (see Table S6 for primers). Families with a unique T-DNA insertion locus were selected on the basis on a clear 1:1 segregation of seed GFP fluorescence in the T1BC1 generation. T1BC1 hybrids, obtained after backcrossing T1 plants with Cvi-0, were screened by PCR for the absence of the T-DNA and selfed. The selection of unmodified Cvi-0 nuclear background (homozygote Cvi-0 at L3) was carried out by genotyping the L3 genomic region in the T2BC1 generation (see Table S6 for primers).

At each generation, the fertility of plants was assessed in the greenhouse by visual examination of silique length, as a proxy for seed production by selfing. Their pollen viability was examined after Alexander staining (Alexander, 1969).

#### Analysis of mutated mitochondrial genomes

Total genomic DNA was extracted from 1.5 g of frozen rosette tissue, pooled from 17-day-old plantlets of the T3BC1 generation grown *in vitro*, using the NucleoSpinPlantII Maxi kit from Macherey Nagel according to the supplier’s instructions. Samples were commercially sequenced using Illumina technology (Eurofins). Reads were aligned with the [Sha]Cvi genome, reconstituted with the Cvi-0 nuclear genome (Simon *et al*., 2022) and Sha organelle genomes (this work). Alignments were visualized with the Integrative Genomics Viewer tool (IGV_ 2.12.3, Thorvaldsdottir *et al*., 2013). PCR amplifications targeting specific regions of the mitochondrial genome were carried out using the primers described in Table S6.

### Estimating the pollen development stage from the bud development stage

Observation of nuclei is the usual procedure to assess the stage of development of pollen in a sample, as pollen mitoses are the milestones of this development (Twell, 2011). However, a different indicator to estimate the stage of pollen development is required when pollen mitoses cannot be directly observed or are perturbed. We exploited the proportional behavior between pollen stage and pistil size that was previously established in Col-0 (Fig. S3). After validation of this correspondence in Cvi-0, we added observations on other floral organs (Table 2) to facilitate the sampling of buds containing the appropriate pollen stage. As pollen development is not synchronized between long and short stamens, the correlation was valid for long stamens only. Hence, only long stamens were sampled for analyses at any specific pollen stage.

**Table 2:** Correspondence between pollen and bud developmental stages in Cvi-0.

| stage of pollen<sup>a</sup> | abbreviation | pistil size (mm) | petal color and length |
| --- | --- | --- | --- |
| tetrad | Td | 0.75 | greenish petals shorter than stamens |
| uninucleate microspore | UNM | 1 | greenish petals shorter than stamens |
| first pollen mitosis | PMI | 1.25 | greenish petals and stamens of same length |
| bicellular pollen | BCP | 1.25-1.5 | greenish petals longer than stamens |
| second pollen mitosis | PMII | 1.5 | greenish petals longer than stamens |
| tricellular pollen | TCP | 1.5-2 | white petals longer than greenish to yellowish stamens |
| mature pollen | MP | 2 | white petals longer than yellow stamens |

<sup>a</sup>in long stamens.

### Analyses of ATP content in plants expressing the ATP ratiometric sensor

#### Generation of plants expressing the ATP FRET biosensor

We cloned the sequence for the MgATP^2-^-specific FRET-based biosensor ATeam1.03-nD/nA (De Col *et al*., 2017) into the pUBDEST vector, *i.e.* under the control of the promoter of *UBQ10*. The construct was introduced into Arabidopsis Cvi-0 by floral dipping. Three out of a total of sixteen T1 individuals that showed biosensor expression as apparent from their YFP fluorescence were chosen and T2 seedlings grown on agar plates were screened for bright fluorescence. One of those lines was used to introduce the biosensor into the [Sha]Cvi and [Kz9]Cvi genotypes by crossing. Hybrid seeds were sown *in vitro*. Six-day-old plantlets per genotype were observed to check the fluorescence and the brighter ones were kept for observation and transferred in the greenhouse until flowering for pollen observation. The long anthers were dissected from buds of the desired stage (Table 2), put in a water drop and gently sectioned with a needle to let pollen grains disperse.

#### Confocal imaging of pollen

Confocal imaging was conducted using a Zeiss LSM780 microscope equipped with a ×40 (C-Apochromat, 1.20 N.A., water immersion) lens as described previously (Wagner *et al*., 2015; De Col *et al*., 2017). ATeam1.03-nD/nA was excited at 458 nm, and the fluorescence signals of mseCFP and cp173-mVenus were captured within the ranges of 475-500 nm and 525-550 nm, respectively, while the pinhole was adjusted to 5.78 airy units. Carbonyl cyanide m-chlorophenyl hydrazone treatment on pollen of fertile plants was used to verify that a depletion in ATP could be detected with lower fluorescence ratios.

#### Ratiometric analyses

Images were processed as established previously (De Col *et al*., 2017) using a custom MATLAB-based software (Fricker, 2016), which involved x-y noise filtering, background subtraction and region-of-interest analysis. Ratio images were calculated on a pixel-by-pixel basis. More precisely, in the case of pollen grains, we set the linear ratio scale between 0.1 and 3.0 with a maximum intensity scale of 250 for the ATeam1.03-nD/nA probes. We corrected for background signal in the measured emission channels by selecting one ROI in each image in the cytosol of a representative non-fluorescent pollen grain. For the ratiometric analysis, 16-pixel large ROIs were selected for all fluorescent pollen grains in one image, which approximately covered the cytosolic and nucleic area, but did not include areas of the intine and exine. The ratiometric data were combined and the obtained measurement dataset was imported within the R software (version 3.6.3) for graphical display and statistical analysis. For each ‘developmental stage’, fixed effect models were used to test the effect of the genotype by comparing models with and without genotype as a fixed factor, with plant/image as nested random factors for analysis. The R package lme4 (Bates *et al*., 2015) was used for fitting random models to the data using Maximum Likelihood estimation. Models were compared by performing a likelihood ratio test.

### Production and analysis of plants expressing the male germline marker

#### Production of sterile and fertile lines expressing the male germline marker

The vector carrying the pTip5;1::H2B-GFP construct (Borg *et al*., 2011) was kindly provided by David Twell, from Leicester University. We introduced it into Arabidopsis Cvi-0 by floral dipping. We crossed [Sha]Cvi and [Kz9]Cvi genotypes as female parents with a homozygous T2 plant for which the GFP fluorescence of nuclei in the male germline was verified in 100% of pollen grains. We used three sterile and two fertile plants and one to three buds per plant for each stage, based on to the bud developmental stage (Table 2). The four long anthers of each bud were dissected and pollen grains were gently dispersed out of the anthers with a needle in a drop of Citifluor AF1 (AgarScientific, Stansted, United Kingdom).

#### Confocal acquisitions and counting

Z-stacks of pollen grains were acquired using a Leica SP8 confocal microscope, using an argon laser line for GFP excitation at 488 nm and detecting fluorescence between 510 and 530 nm. Pollen grains with no, one or two fluorescent nuclei were counted to estimate the proportion of pollen grains expressing the male germline marker. Pollen grains with abnormal shape were assumed to be dead and were not taken into account.

### Monitoring of mitochondria morphology in pollen

*Construction of the pMDC32-UBX1 vector for pollen specific expression of transgenes*.

We constructed a vector aiming for specific expression in pollen using the promoter of the *Brassica napus UBX1* gene, shown to be specifically active in the male gametophyte (Gallois *et al*., 2013). We amplified a 603 bp fragment containing the promoter and the 5’ UTR of the *BnUBX1* gene from genomic DNA of the Brutor cultivar with primers adding *Hin*dIII and *Kpn*I restriction sites (Table S6). This fragment was cloned into the pMDC32 vector (Curtis and Grossniklaus, 2003) in place of the *35S* promoter. The resulting vector, named pMDC32-UBX1 was used to express the matrix mitochondria-targeted GFP, mt-GFP (Logan and Leaver, 2000) in Cvi-0 plants by floral dipping. Expression of the mt-GFP gene was verified for all developmental stages of pollen in T1 plants using fluorescence microscopy (Fig. S4).

#### Generation of plant lines homozygous for a pollen mt-mCherry and the male germline GFP markers

We cloned the coding sequence of the mCherry fluorescent protein, fused to the same mitochondrial targeting sequence as above, into the pMDC32-UBX1 vector and transformed Cvi-0 plants. A homozygous Cvi-0 T2 was crossed on a [Kz-9]Cvi plant carrying the pTip5;1:H2B-GFP construct. After selfing, a plant homozygous for both markers was selected on the basis of mCherry and GFP fluorescence in all the pollen grains. This genotype was used to cross [Sha]Cvi, and the cross was repeated on the F1. Plants from the second generation were test-crossed with Col-0 and back-crossed again with the [Kz-9]Cvi doubly marked line. Two [Sha]Cvi families homozygous for both markers were identified on the basis of 100 % transmission of both markers, tested by PCR, to the progenies of the test-cross (see Table S6 for primers).

#### Confocal acquisitions

Buds from 4- to 6-week-old plants were dissected to extract the long anthers at the desired stage of development (Table 2). Pollen grains were dispersed out of the anthers with a needle in a drop of Citifluor AF1 (AgarScientific, Stansted, United Kingdom). Confocal imaging was conducted on a Leica SP8 microscope equipped with a 40x (Leica HC PL APO CS2, 1.30 numerical aperture, oil immersion) lens. Samples were illuminated at 561 nm for mCherry excitation and at 488 nm for GFP. The mCherry fluorescence signal was captured in the 590-630 nm range, while GFP was captured in the 498-520 nm range. Z-stacks of pollen grains were acquired in bidirectional mode at 700 Hz, with the pinhole adjusted at 0.5 airy unit, voxel size set at 75 nm x 75 nm x 230 nm (xyz) and grayscale resolution set at 16-bits.

#### Image analysis

The channels in the acquired TIF images were split and saved in distinct images. The autofluorescence of pollen wall in the GFP channel was used for the segmentation of pollen grains. All operations for segmenting pollen grains and mitochondria from the GFP and mCherry fluorescence channels were performed using the Biological Image Processing (BIP) software (freely available at https://free-d.versailles.inrae.fr/html/bip.html). The BIP pipeline operator was used to apply at once the segmentation workflows to the complete image dataset.

Images with autofluorescence signal were resampled into cubic voxels and downscaled by a factor of 0.3 along each dimension. Anisotropic diffusion followed by a median filter was applied to filter noise and improve signals in the images, while preserving contrasts at grain boundaries. A Gaussian gradient operator was then applied to highlight contour positions in the pollen grains. To avoid over-segmentation, non-significant minima in the resulting image were filtered out before running the watershed transform. Labels touching the border of the images, typically corresponding to grains that were not completely included within the 3D field-of-view, were automatically removed. Post-processing steps were applied to remove small objects, fill holes inside objects, and regularize object contours. Merged objects were separated by running the watershed transform on an inverted distance map between object and background voxels. The segmented grains were finally crop and their individual masks stored separately in different images. The complete pipeline for segmenting grains is available in the *segment-grains.pipeline* file that has been deposited at <https://doi.org/10.57745/56PALB>. The individual cropped images of the GFP channel were used to visually assess the presence of the generative nucleus in each analysed grain of the ‘bicellular pollen’ stage.

For each grain, the same crop as above was applied to the channel with mCherry fluorescence intensity. A difference-of-Gaussian filter was applied to enhance signals at mitochondria. Automated binarization was performed using an in-house algorithm, where the selected threshold corresponds to the maximization of object size homogeneity. Touching mitochondria were split using the procedure described above for separating merged pollen grains. The binary mask of the pollen grain was used to selectively retain the detected mitochondria in that grain. The complete pipeline for segmenting mitochondria is available in the *segment-mitochondria.pipeline* file at <https://doi.org/10.57745/56PALB>. Object measurements were performed using the image analysis operator available in BIP. The volume of each segmented mitochondrion was estimated by multiplying its number of voxels by the individual voxel volume. The obtained measurement table was imported into the R software (version 3.6.3) for graphical display and statistical analysis. For each stage, mixed effect models were used to test the effect of the genotype by comparing models with and without genotype as a fixed factor, with date/bud as nested random factors. The R package lme4 (Bates *et al*., 2015) was used for fitting models to the data using Maximum Likelihood estimation. Models were compared by performing a likelihood ratio test.

## Results

### *orf117Sha* is the causal gene for Sha CMS

The fertility phenotype of the [Kz9]Cvi genotype (Table 1), which is indistinguishable from that of Cvi-0, confirmed that the Kz-9 cytoplasm is not a CMS inducer (Fig. 1). In a previous work, we identified *orf117Sha* as a candidate gene causing cytoplasmic male sterility by comparing the mitochondrial genomes of Sha and Kz-9 (Gobron *et al*., 2013). Intriguingly, its RNA expression profile was not affected in restored genotypes (Durand *et al*., 2021). Since our initial comparison of Sha and Kz-9 mitochondrial genomes did not completely cover the mitochondrial genome sequence, we could not exclude the possibility that the causal gene for CMS remained undetected. To address this shortcoming, we sequenced, *de novo* assembled and annotated the organellar genomes of these two genotypes. Their chloroplast genomes are almost identical (Accession numbers: Kz-9, OY747152, Sha, OY747153; Fig. S5). The Sha and Kz-9 mitochondrial genomes (Accession numbers: OY747154 and OY747151, respectively; Fig. S6) are remarkably collinear (Fig. 2A). The major difference lies within a small region (3.5 kb in Kz-9, 6 kb in Sha) that is not shared between the two genomes; in Sha, this specific region carries *orf117Sha* (Fig. 2B). We did not detect any other open reading frame (orf) of at least 100 codons specific to the Sha mitochondrial genome compared to Kz-9 and Col-0 (Table S9). Therefore, *orf117Sha* remained the best candidate for causing Sha-CMS.

**Fig. 1.**
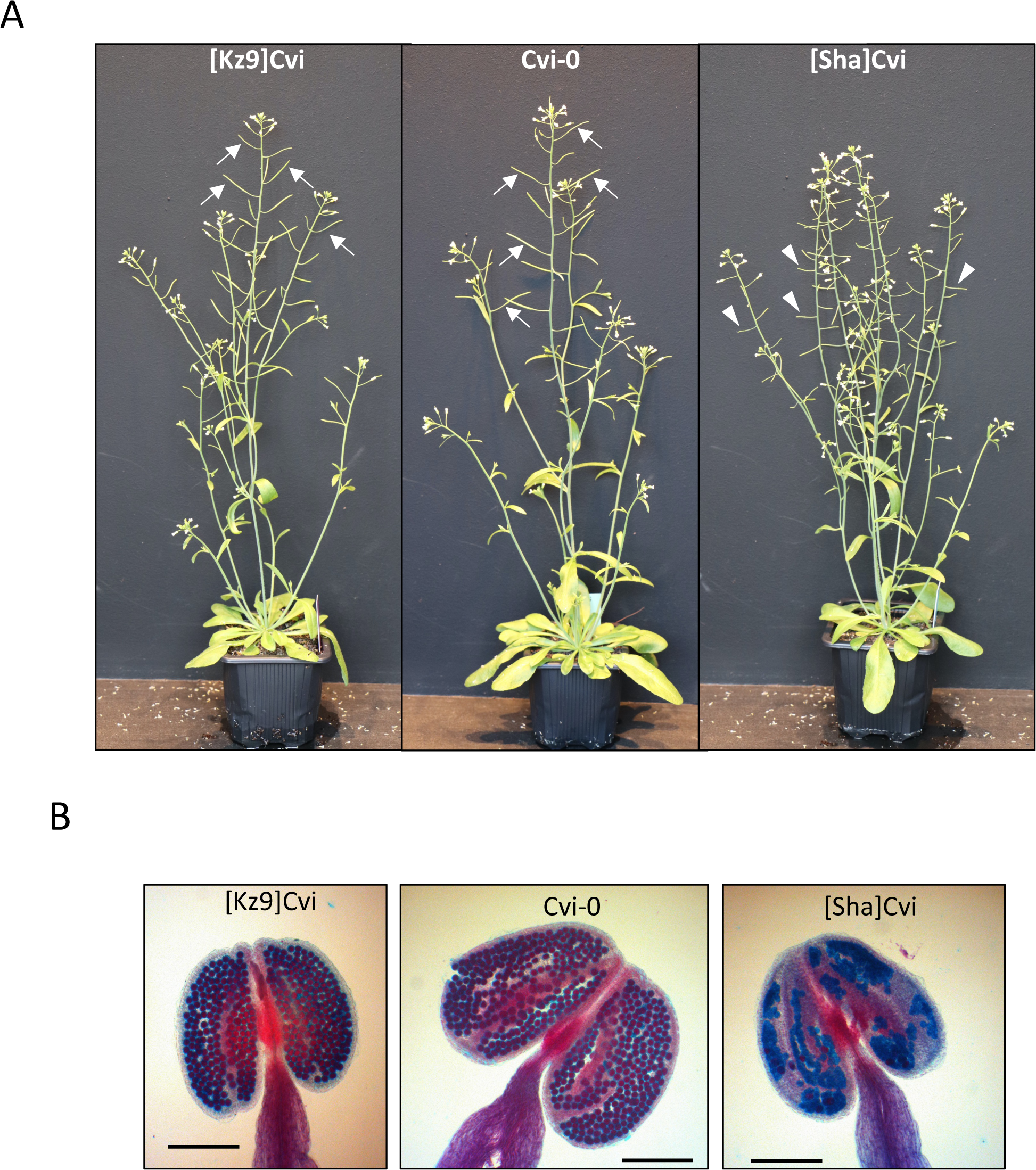
[Kz9]Cvi and Cvi-0 have undistinguishable pollen fertility phenotypes. A: [Kz9]Cvi, Cvi-0 and [Sha]Cvi plants 5 weeks after sowing. Arrows: siliques producing seeds; arrowheads: empty siliques. B: Alexander staining of pollen of the plants shown in A. Viable pollen grains are stained in red; dead ones appear in blue; scale bar: 200 µm.

**Fig. 2.**
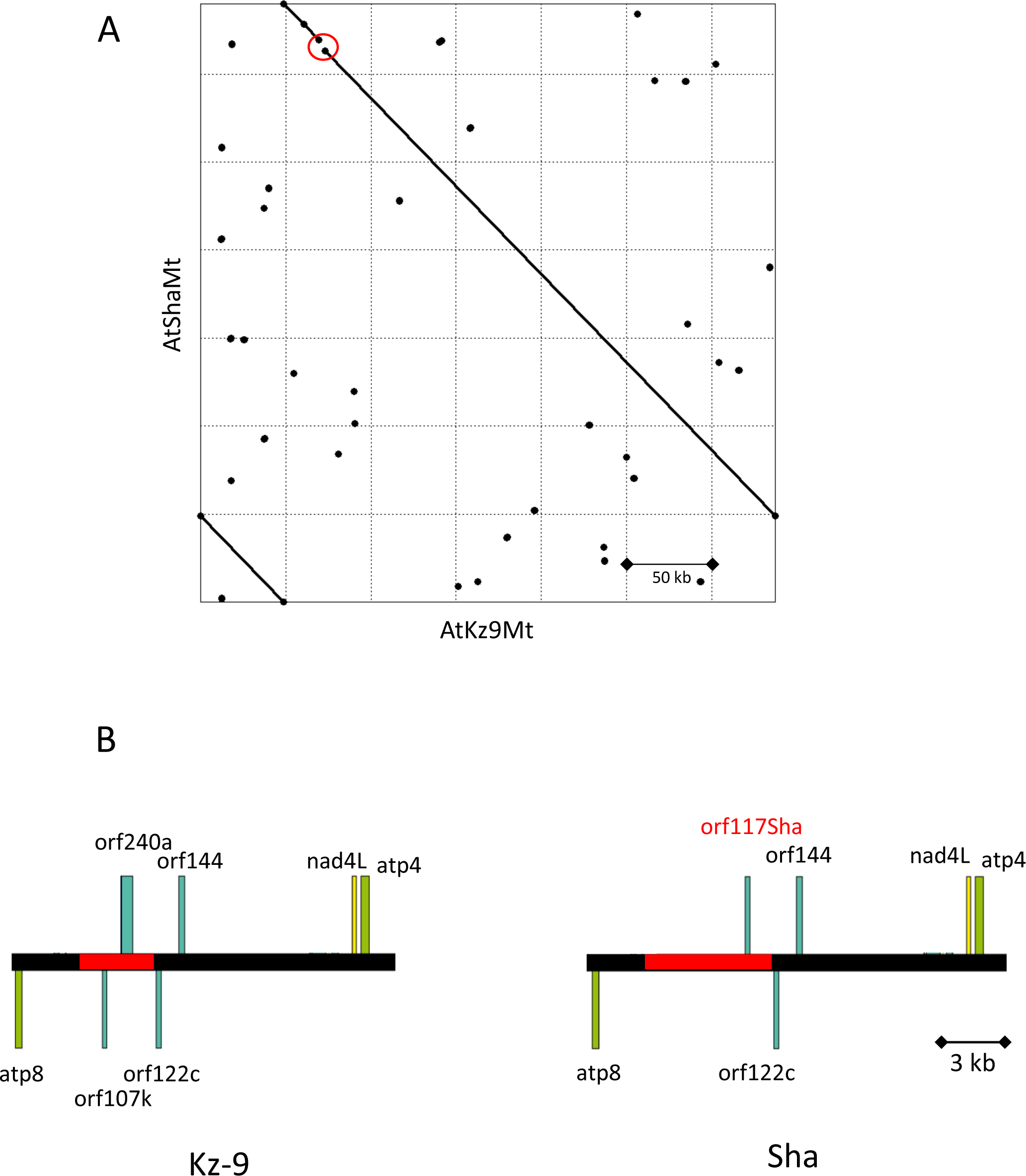
Kz-9 and Sha mitochondrial genomes are almost identical. A: Dotplot alignment of Kz-9 and Sha mitochondrial sequences. The sequences have been assembled in opposite orientation. The main structural variation (red circle) corresponds to the *orf117Sha* region shown in B. B: Close up of the regions unshared by the two genotypes. The extent of specific sequence for each genotype is highlighted in red. Annotation has been simplified for clarity. Drawn with OGDraw.

As the ORF117SHA sequence is similar to that of ORF108 (Gobron *et al*., 2013), which has been reported to induce sterility in Arabidopsis when produced in transgenic plants (Kumar *et al*., 2012), we attempted to phenocopy the CMS phenotype by transgenesis. The coding sequence of *orf117Sha* was fused to a mitochondrial signal peptide, placed under the control of the UBQ10 promoter, which is active in pollen (Fig. S1), and introduced into Cvi-0. As Sha-CMS is gametophytic (Simon *et al*., 2016), we expected ORF117SHA to cause 50% pollen death in T1 plants, and lead to 50% of transgenic plants in the T2 generation. We analyzed pollen viability in 21 T1 plants: only one had approximately 50% aborted pollen, one was completely sterile, and five additional T1s had between 25% and 50% dead pollen. We analyzed the transmission of the T-DNA to the selfing progenies of these T1s, except for the sterile one, and in four additional T2s whose mothers had few or no dead pollen (Table 3). The progeny of the unique T1 with ∼50% dead pollen displayed a strong segregation bias, with almost no transmission of the transgene (four positives out of 306). Three other T2 plants presented a mild but significant deficit in plants carrying the transgene, but with no apparent correlation with the amount of dead pollen in the mother plant (Table 3). While these observations are compatible with a role of *orf117Sha* in inducing male sterility, they were insufficient to reliably establish such a role.

**Table 3:**
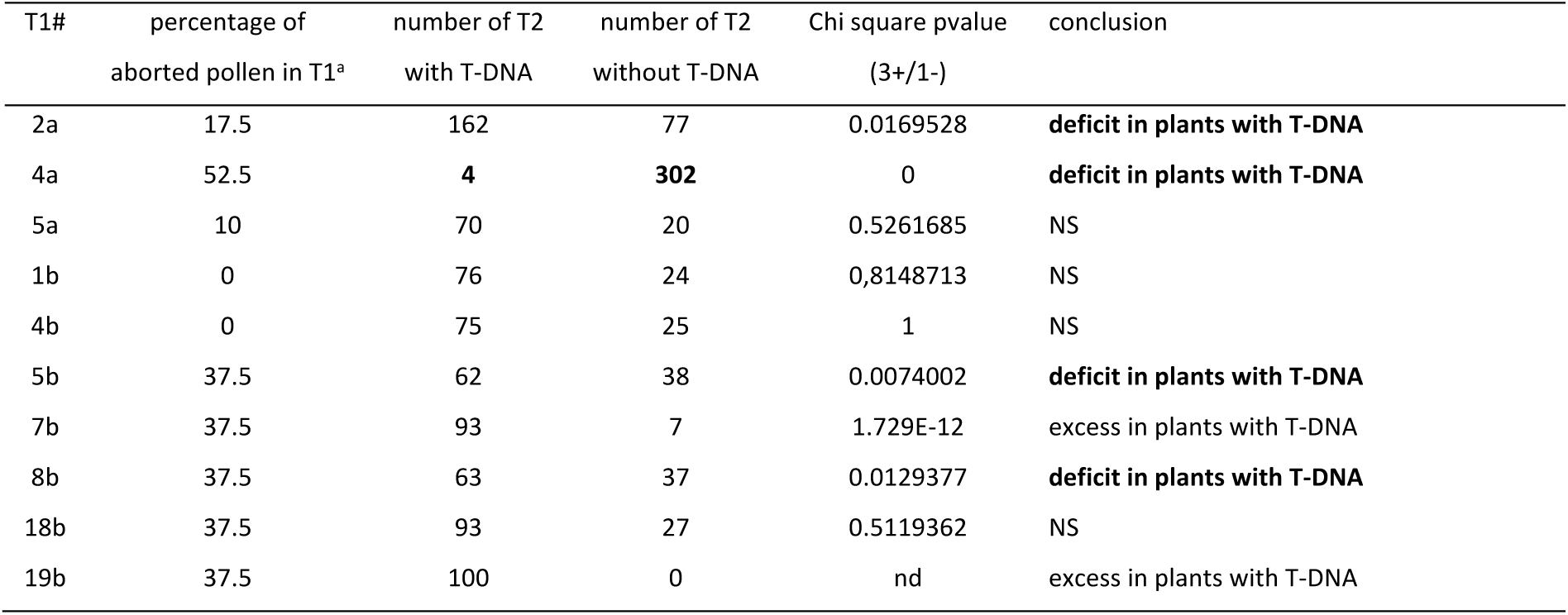
Analysis of transgene transmission in selected T1s with the *orf117Sha* construct.

Seeking decisive evidence, we targeted *orf117Sha* using mitoTALENs. This strategy has recently been successfully used to reverse CMS phenotypes to fertility in rapeseed and rice by inducing deletions of the mitochondrial CMS genes through the repair of mitoTALEN-induced DNA breaks (Kazama *et al*., 2019). We used two pairs of mitoTALENs, designated mTAL1 and mTAL2 (Fig. S2), to target *orf117Sha* in [Sha]CviL3^S^ plants (Table 1). These plants are partially restored because they possess a weak restoring Sha allele on chromosome 3 (Durand *et al*., 2021). They are clearly less fertile than Cvi-0 but produce enough selfing seeds for floral dipping transformation (Fig. 3A, B). We selected ten and eight independent transformants for mTAL1 and mTAL2 constructs, respectively. T1 plants obtained with mTAL1 were as fertile as Cvi-0 (*e.g.* mTAL1#2), except one (mTAL1#1), which displayed a phenotype similar to that of [Sha]CviL3^S^ (Fig. 3 C, D). Conversely, T1 plants obtained with mTAL2 were only partially fertile (*e.g.* mTAL2#5), except for one (mTAL2#2) that was as fertile as Cvi-0 (Fig. 3 C, D). To obtain plants with stable mutated mitochondrial genomes in a Cvi-0 nuclear background, we backcrossed all T1s with Cvi-0. For further analysis, we selected T1BC1s without the mitoTALEN construct from T1s with single T-DNA insertions (Fig. 4A). All T1BC1 plants from mTAL1 were fertile, whereas most T1BC1 plants from mTAL2 were sterile (Table S10). PCR amplification showed the presence of *orf117Sha* in sterile or partially fertile plants, whereas it was not detected in fertile plants, excepted one (mTAL2#2- 3, Table S10). This suggests that the fertility phenotype of the plants was linked to the mitoTALEN-induced loss of *orf117Sha*. After selfing of T1BC1 plants, we selected and observed 28 T2BC1 plants that were homozygous for the Cvi-0 allele at the bottom of chromosome 3, thus possessing a Cvi-0 nuclear background (Fig. 4A). All were fertile, except for two partially sterile plants from the mTAL1#8- 3 family (Table S11). We produced genome-wide Illumina sequences from four T2BC1 plants from distinct T1BC1 families (Fig. 4B, C) and the [Sha]Cvi control. Alignments of paired reads on the genome of [Sha]Cvi revealed 2 to 12 kb deletions that included both *orf117Sha* and its neighbor *orf122c* in the mitochondrial genome of the four mutants (Fig. 5A). *orf117Sha* and *orf122c* were undetectable or detected at very low levels by PCR in all T2BC1 plants (Table S11, Fig. S7), indicating that both genes were effectively removed from fertile mitochondrial mutants. Although different in the four samples, the deletions shared the same left border, which was due to recombination between the copies of the 1 kb repeat (R_Sha_2) present upstream of *orf117Sha* and *cob* in Sha (Fig. 5B, C) and absent from Kz- 9 (Table S5). This was also the case in all fertile T2BC1s (Fig. S7 and Table S11). Interestingly, the *rpl5-cob* region was also detected in fertile plants (Fig. 5C), indicating a duplication of the *cob* gene in these plants, which is supported by the raise in coverage of Illumina reads for this gene and upstream sequences (Fig. S8). By examining the clipped sequences of reads at the right borders of the deletions, we found very short sequences (<50 bp) that were repeated elsewhere in the mitochondrial genome (Table 4). We showed that these microhomologies had been used to repair the mitoTALEN DNA breaks by detecting the expected recombined sequences by PCR (Fig. S9A) and we observed an abrupt increase in read coverage at these positions in fertile samples (Fig. S9B), as for *cob*.

**Fig. 3.**
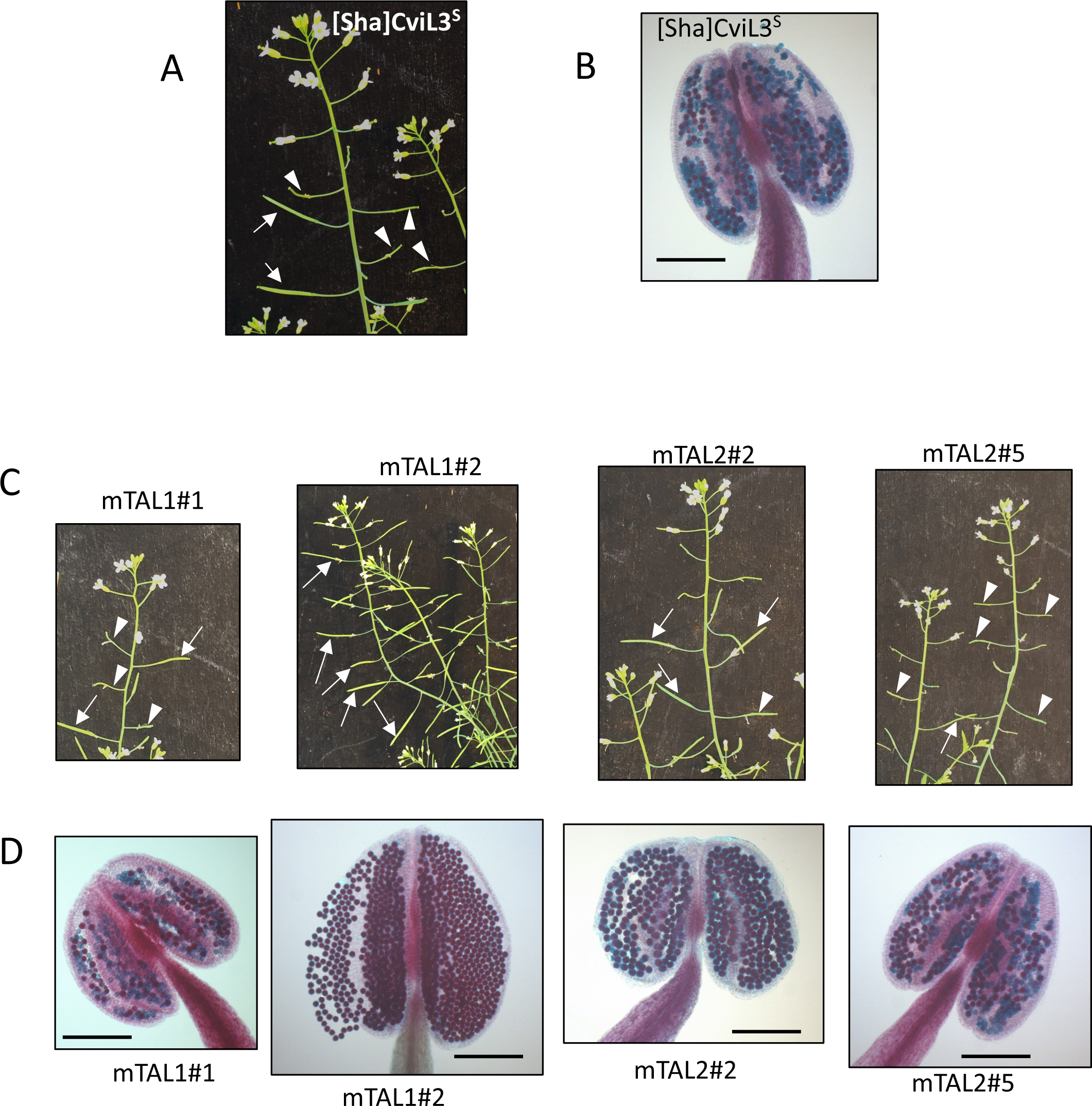
Fertility phenotypes of the recipient genotype and mitoTALEN T1 transformants. A, B. Fertility phenotypes of [Sha]CviL3^S^, the recipient genotype. C, D. Phenotypes of T1 plants. mTAL1#1: the only plant with the mTAL1 construct that was not fully fertile, mTAL1#2: representative T1 plant with mTAL1 construct, mTAL2#2: the most fertile plant with the mTAL2 construct, mTAL2#5: representative plant with the mTAL2 construct. A, C: Global seed fertility at the plant level. Arrows: siliques producing seeds; arrowheads: empty siliques. B, D: Alexander staining of pollen. Viable pollen grains are stained in red; dead ones appear in blue; scale bar: 200 µm.

**Fig. 4.**
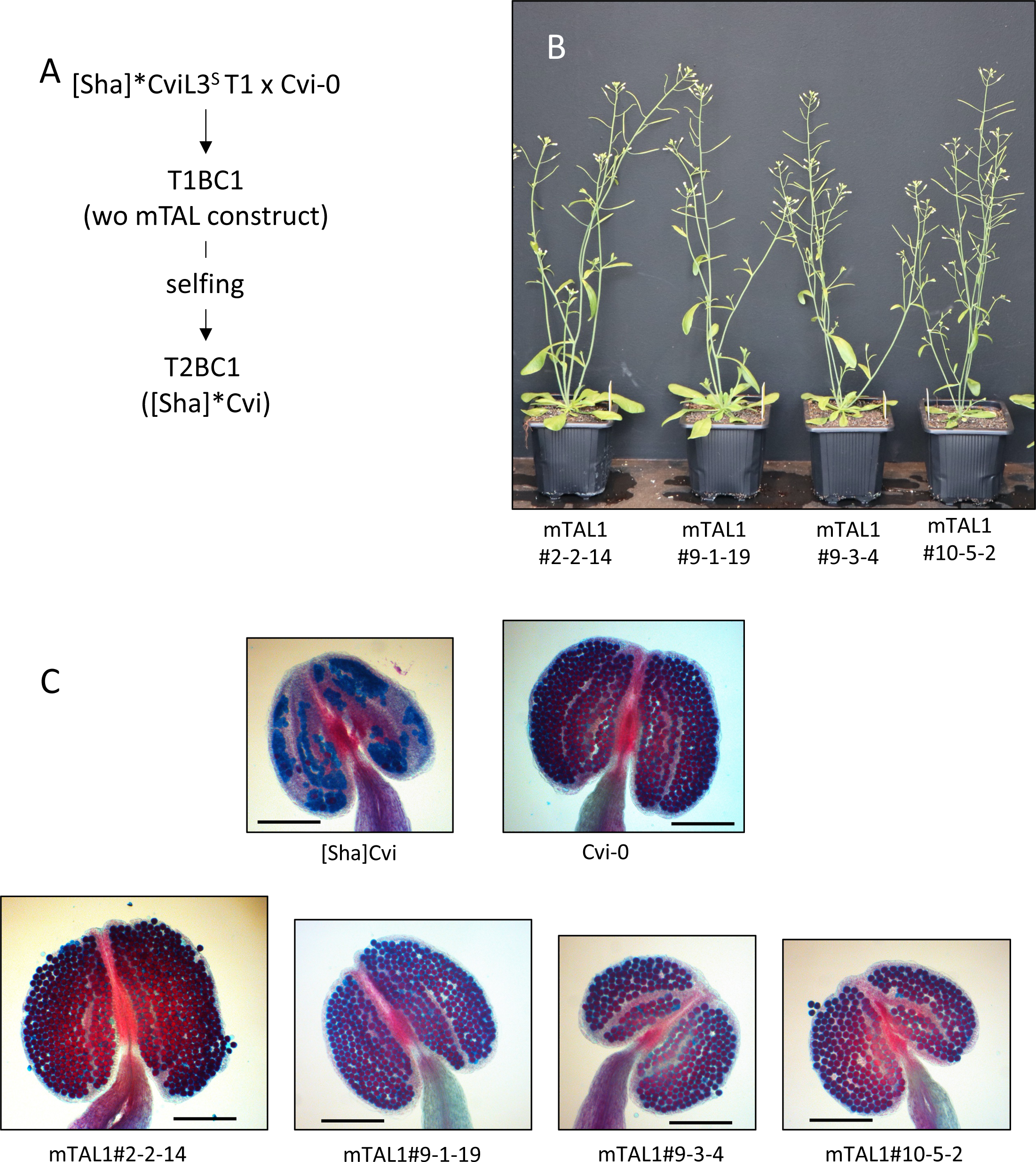
Analysis of revertant mitochondrial mutants in the Cvi-0 nuclear background. A. Recovery of mutated mitochondrial genomes in the Cvi-0 nuclear background without mitoTALEN construct. The * indicates that the Sha mitochondrial genome has been modified by mitoTALEN. B, C. fertility phenotypes of T2BC1 plants selected for genome wide sequencing. B. global plant phenotypes. C. Alexander stainings of pollen. [Sha]Cvi and Cvi-0 are shown as sterile and fertile controls. Viable pollen grains are stained in red; dead ones appear in blue; scale bar: 200 µm.

**Fig. 5.**
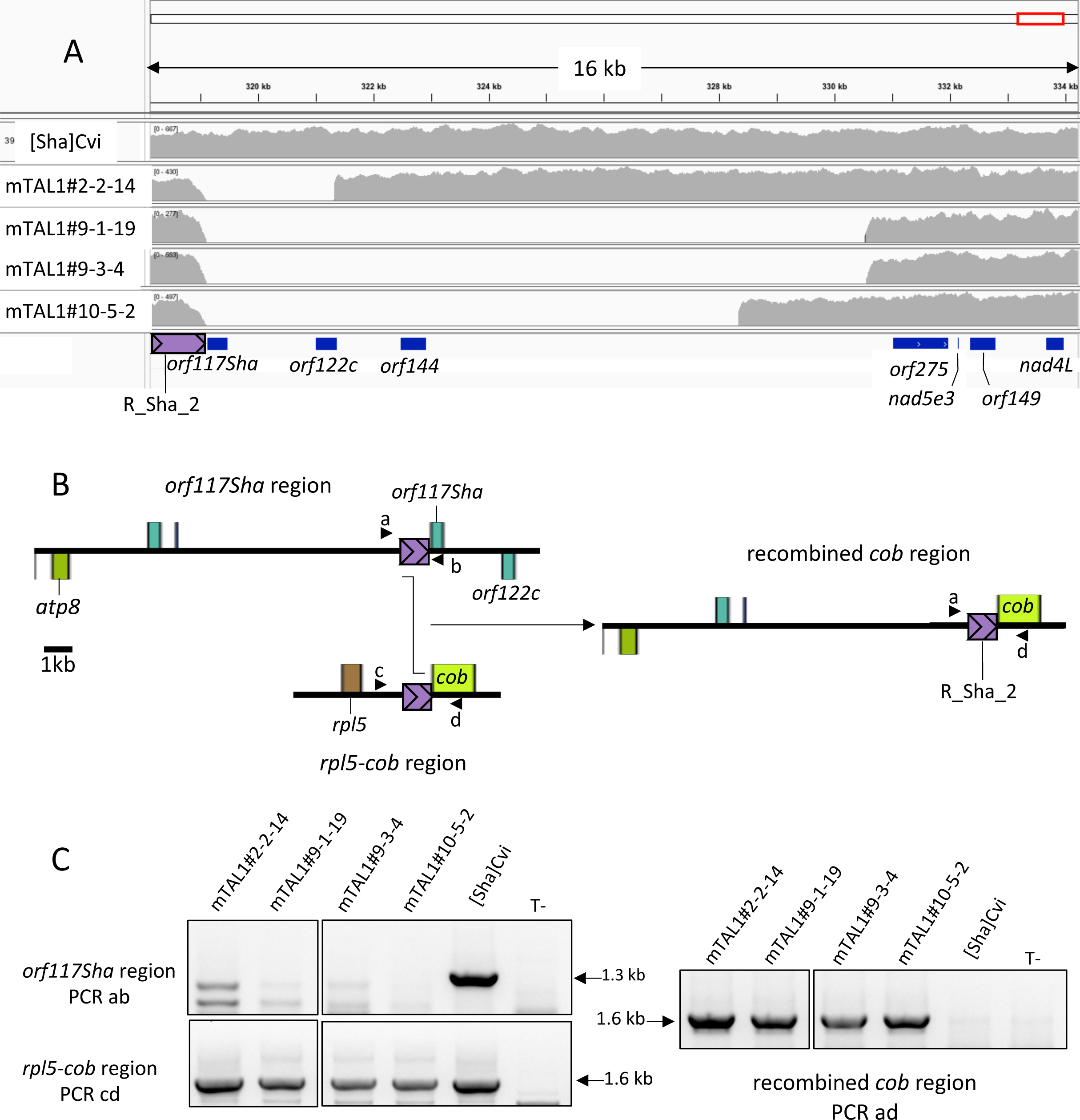
Repair of mitoTALEN-induced DNA breaks in mitochondrial genomes of fertile plants. A. IGV visualisation of read coverage for the *orf117Sha* region for [Sha]Cvi (sterile control) and the four sequenced mitochondrial mutants. B. Regions carrying the R_Sha_2 repeat. *orf117Sha* and *rpl5-cob* regions are present in the Sha mitochondrial genome and may recombine to give the recombined *cob* region; arrowheads (a, b, c and d) indicate the positions of primers used for PCR (Table S6). C. PCR detection of the three environments of R_Sha_2 shown in B; non relevant lanes of the gel have been removed for clarity. T-: no DNA. The expected amplification sizes are indicated on the side.

**Table 4:** Very short repeats (VSR) used for repair of mitoTALEN induced DNA breaks.

| repeat number | recombined in mutant | length of repeat | orientation of copies in the Sha genome | position downstream of *orf117Sha* | second position in Sha mitochondrial genome |
| --- | --- | --- | --- | --- | --- |
| VSR-1 | mT1#2-2-14 | 13 bp | inverted | 321,344-321,356 | 124,125-124,137 |
| VSR-2 | mT1#9-1-19 | 16 bp | direct | 330,531-330,546 | 163,711-163,726 |
| VSR-3 | mT1#9-3-4 | 47 bp | direct | 330,564-330,610 | 92,237-92,283 |
| VSR-4 | mT1#10-5-2 | 29 bp | inverted | 328,353-328,381 | 56,793-56,821 |

Considering the above results, the deletion of *orf117Sha* and *orf122c* is the only possible cause of reversion to fertility in mitoTALEN-induced mutants. As *orf122c* is present not only in Sha but also in Kz-9 and Col-0 (Table S9), which are not CMS-inducers, we conclude that *orf117Sha* is the causal gene for Sha-CMS.

### Pollen abortion process in the Sha CMS

We further investigated Sha-CMS by implementing pollen-centered approaches. From our analysis of Sha and Kz-9 organelle genomes, we selected [Kz9]Cvi as the best fertile control to observe events associated with *orf117Sha*-induced pollen abortion. In order to use equivalent developmental stages in both genotypes for comparison, we used the bud developmental stage as a proxy to the pollen developmental stage (Table 2). For clarity, the developmental stage will therein be between ‘quotes’ when it refers to the stage inferred from the bud developmental stage.

#### Pollen abortion is not accompanied by a deficit in ATP

ATP limitation has often been assumed to be the immediate cause of pollen abortion in CMS (Hanson and Bentolila, 2004; Chase, 2007; Yang *et al*., 2022), including in the gametophytic HL-CMS of rice (Wang *et al*., 2013). However, this hypothesis has been questioned (Touzet and Meyer, 2014). We harnessed the cytosolic ATeam1.03nD/nA protein (hereafter called cATeam), a fluorescent FRET-based biosensor for Mg-ATP^2-^ concentration (De Col *et al*., 2017), in order to experimentally address *in vivo* whether a deficit in ATP could be the trigger of, or take part in, pollen abortion in the Sha-CMS. We transformed Cvi-0 with the cATeam coding sequence under the control of the *UBQ10* promoter and introduced the construct into the [Kz9]Cvi and [Sha]Cvi genotypes by crossing. We acquired images of pollen at the ‘bicellular pollen’ stage and around the second pollen mitosis (‘PMII’), based on bud developmental stages (Table 2). The ratiometric analyses showed no significant difference between the sterile and the fertile at both stages (Fig. 6). As more than 80% of the pollen is already dead in the sterile at the ‘PMII’ stage (Durand *et al*., 2021), it is likely that the abortion process was already engaged in the rare pollen grains which could be imaged for this stage. Therefore, a deficit in ATP content is very unlikely to trigger or participate in pollen abortion in [Sha]Cvi sterile plants, thus suggesting a different, still unveiled, physiological cause for the triggering of pollen abortion. We then focused on the events that occur in pollen during the abortion process.

**Figure 6.**
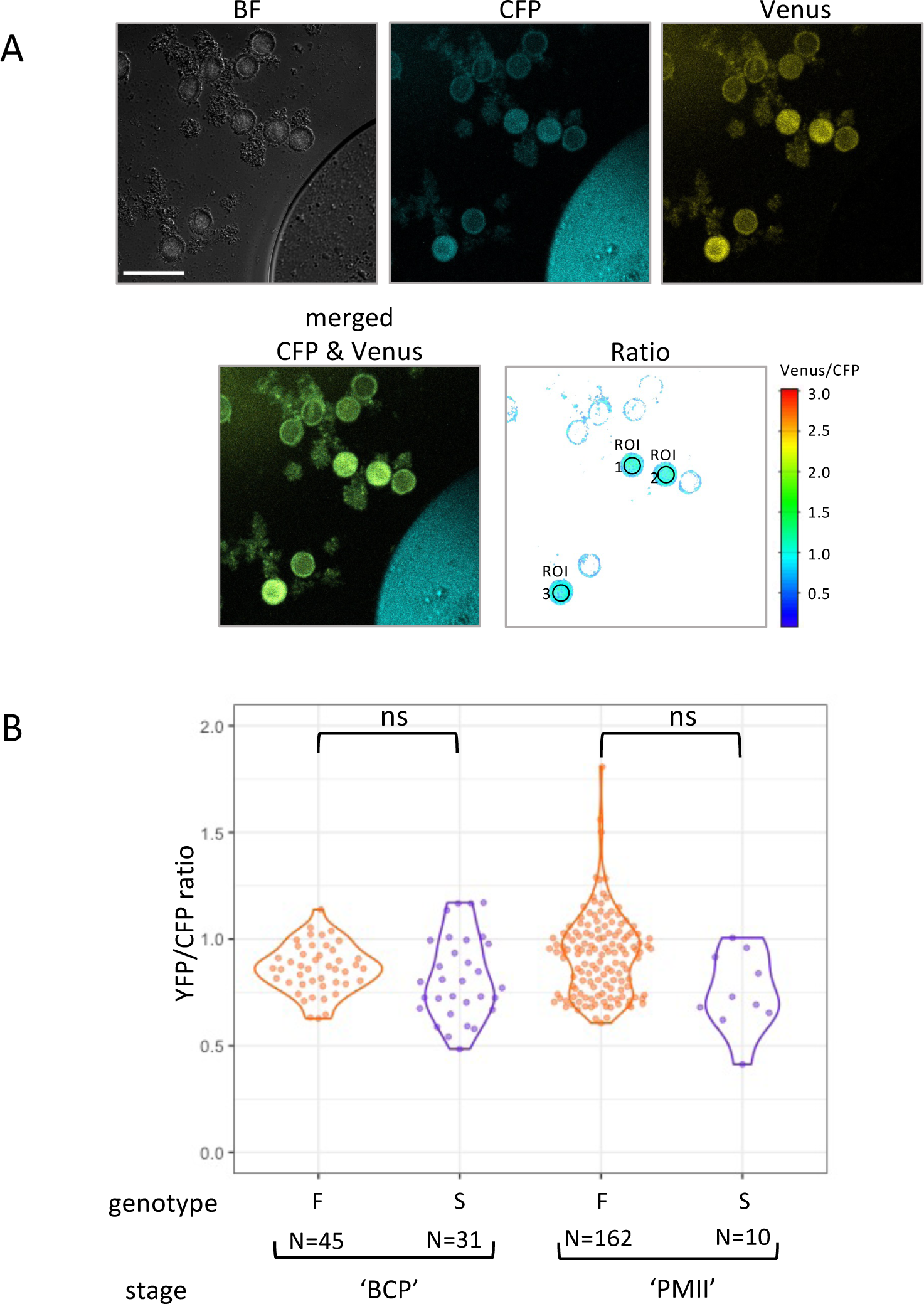
*In vivo* ATP sensing in pollen. A. Representative micrograph of pollen grains in the ‘bicellular pollen’ stage of the fertile genotype expressing the FRET sensor ATeam1.03nD/nA in their cytosol. Note that the plant is heterozygous for the biosensor, resulting in grains showing sensor fluorescence (3 shown here) and grains with background fluorescence only. Brightfield (BF, grey), CFP (turquoise) and Venus (yellow) channels of CLSM images are shown above the merge CFP and Venus image and the ratiometric image of Venus/CFP (rainbow scale) after background subtraction and pixel-by-pixel ratio calculation. Depicted regions of interest (ROIs) were used for the measurement of FRET ratios as an indicator for pollen MgATP content. Scalebar: 50 µm. B. Measures on pollen at the ‘bicellular pollen’ stage (‘BCP’) and around pollen mitosis II (‘PMII’) based on the bud developmental stage (Table 2). At least three buds were dissected for each stage. Each measurement point corresponds to an individual pollen grain. The complete dataset is listed in Table S7. The small sample size of sterile measurements for PMII is due to the scarcity of viable pollen with fluorescence at this stage. F: [Kz9]Cvi fertile genotype, S: [Sha]Cvi sterile genotype, N = number of measurement points. Statistical comparison between genotypes was performed for each stage by comparing mixed linear models with or without considering the genotype as a variable. ns: pvalue of the likelihood ratio test between models > 0.1.

#### The pollen development is desynchronized and prematurely stopped in sterile plants

We previously reported that pollen abortion in [Sha]Cvi occurred during the bicellular stage of pollen development, from ∼ 90 % of viable pollen just after the first pollen mitosis (PMI) to ∼ 2% after PMII (Durand *et al*., 2021). As the PMI coincides with the establishment of the male germline (Twell, 2011), we used the pTIP5;1::H2B-GFP construct, which specifically targets GFP to the nuclei of germinative and sperm cells (Borg *et al*., 2011), to track the male germline and follow the progress of the pollen developmental program in sterile plants compared to fertile ones. We introduced this male germline marker into [Kz9]Cvi and [Sha]Cvi by crossing both genotypes with a homozygous Cvi-0 plant homozygous for the construct. In the resulting hybrids, half pollen grains are expected to carry and express the male germline marker. We counted the proportion of pollen grains with GFP-marked nuclei around PMI, at the ‘bicellular pollen’ stage and around PMII, using the developmental stage of the bud as a proxy to the pollen expected developmental stage (Table 2). This proportion was consistently lower in sterile [Sha]Cvi F1 plants compared to fertile [Kz9]Cvi F1 plants, which presented approximately the expected proportion of GFP marked nuclei at both stages (Fig. 7). This indicated that, in sterile plants, some pollen grains carrying the male germline marker did not express it: either they did not enter into PMI, or they were already dead. Because they tend to agglomerate and could not be numbered, shrunken dead pollen grains were not taken into account, but dead grains are likely to lose the marker fluorescence before shrinking. However, because the proportion of dead pollen at the ‘PMI’ stage was very low, we concluded that, in the sterile, a substantial proportion of the grains that had no GFP-marked nucleus were alive but had not established the male germline. Remarkably, in the following stages, the proportion of grains with one GFP-marked nucleus in the sterile did not significantly change (Fig. 7). This suggests that approximately the same proportion of grains with and without a generative cell died between the examined bud stages. It is also possible that more grains with a generative cell than without one died between the ‘PMI’ and ‘bicellular pollen’ stages, and were compensated by grains which executed PMI during the same period. At the ‘PMII’ stage, when more than 80% of the grains are dead, very few grains had two GFP-marked sperm nuclei in the sterile, whereas in the fertile their proportion was close to the expected 50%. In addition, in sterile plants, grains with two GFP-marked spermatic nuclei co-existed with grains with one GFP-marked generative nucleus, which was never observed in fertile plants (Fig. 7). Therefore, the pollen developmental program was desynchronized in sterile plants. This resulted in the loss of the correspondence between bud and pollen developments for a large proportion of the grains. In addition, our results strongly suggest that pollen grains aborted at variable pollen developmental stages, from before PMI to after PMII.

**Figure 7:**
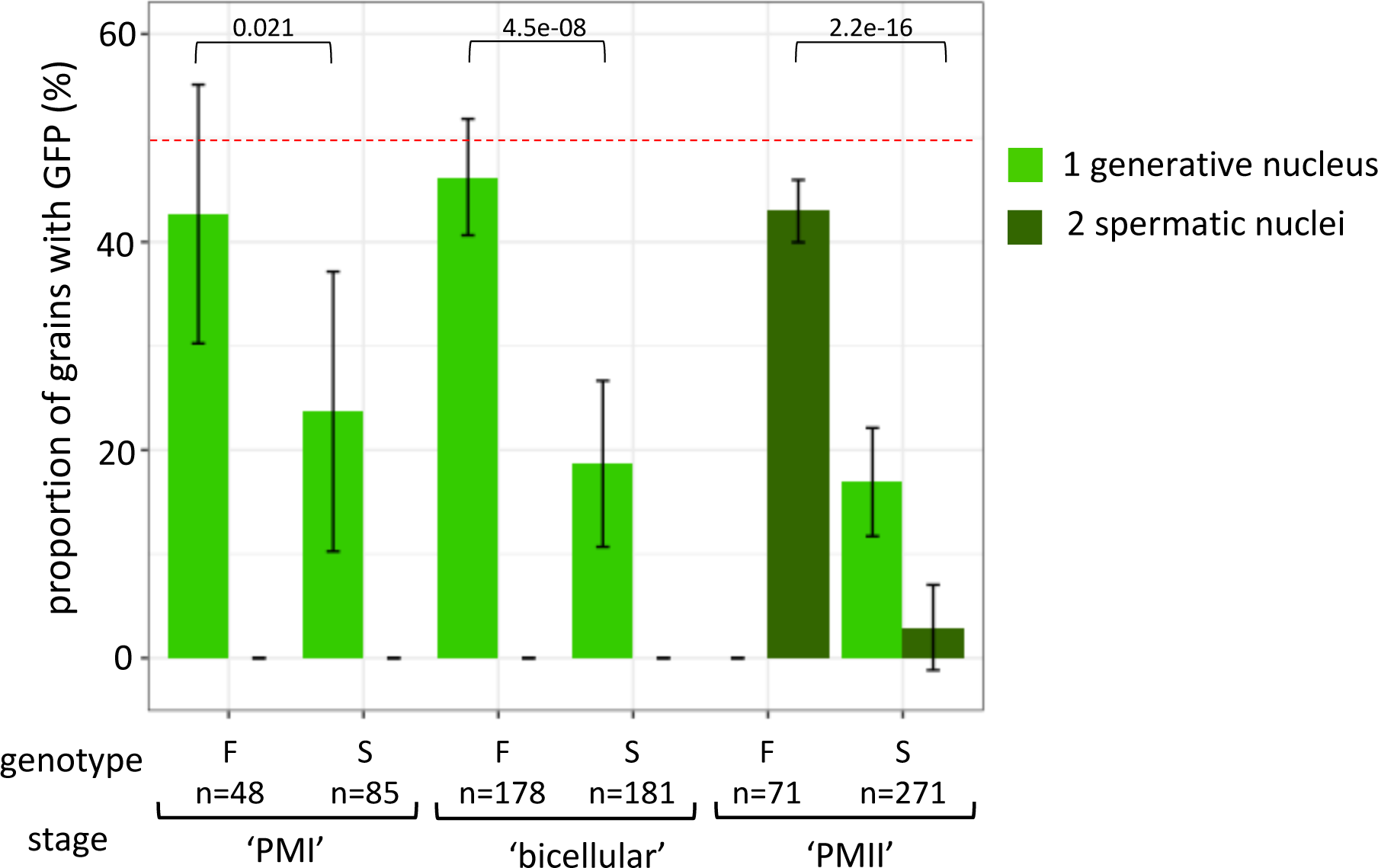
Tracking of male germline during pollen development. Proportions of pollen grains showing GFP fluorescence in one nucleus (generative cell) or two nuclei (spermatic cells) are plotted. Shrunken dead pollen grains were not taken into account in the sterile. F: [Kz9]Cvi, S: [Sha]Cvi, both genotypes heterozygous for the pTIP5;1::H2B-GFP construct. The red dotted line indicates the expected value for plants heterozygous for the male germline marker (50%). Error bars indicate the mean +/- standard deviation amongst buds. Total number of observed grains are indicated under the genotype for each stage. Pollen stages were estimated using bud developmental stage (Table2). The pvalue of Chi square tests performed on the numbers of grains with marked and unmarked nucleus for the ‘PMI’ and ‘bicellular’ stages, and on the numbers of grains with two and less than two marked nuclei for the ‘PMII’ stage are indicated above the bars.

#### Pollen mitochondria swell during the pollen death process

As the genetic cause of pollen abortion resides in mitochondria, we observed mitochondria and pollen germline nuclei in pollen grains of [Kz9]Cvi and [Sha]Cvi plants homozygous for both pBnUBX1::mt-mCherry for mitochondria monitoring and pTIP5;1::H2B-GFP for detection of germline nuclei (Fig. 8A). We imaged isolated pollen grains of the ‘uninucleate microspore’ and ‘bicellular pollen’ stages and measured the average volume of mitochondria for each pollen grain. The results showed that mitochondria of sterile plants were slightly but significantly bigger than those of fertile ones at the ‘uninucleate microspore’ stage, with similar distributions in the two genotypes (Fig. 8B). In contrast, at the ‘bicellular pollen’ stage, pollen grains from the sterile genotype presented a bimodal distribution of the average mitochondrial volume per grain, indicating a swelling of mitochondria in a proportion of the grains (Fig. 8C). As mitochondria swelling has been associated with cell death (Scott and Logan, 2008), we hypothesized that the grains presenting swollen mitochondria were those engaged in the abortion process at the time of observation, which was consistent with the timing of pollen death. In order to address whether mitochondrial swelling was correlated with the establishment of the male germline, we plotted the average volume of mitochondria according to the detection of the generative nucleus thanks to the GFP fluorescence of the male germline marker. The result clearly showed that the swelling of mitochondria was independent from the establishment of the male germline in pollen of the sterile genotype (Fig. 8D). This observation supports that the timing of mitochondria swelling, hence pollen death, is not dependent on the grain developmental stage.

**Figure 8.**
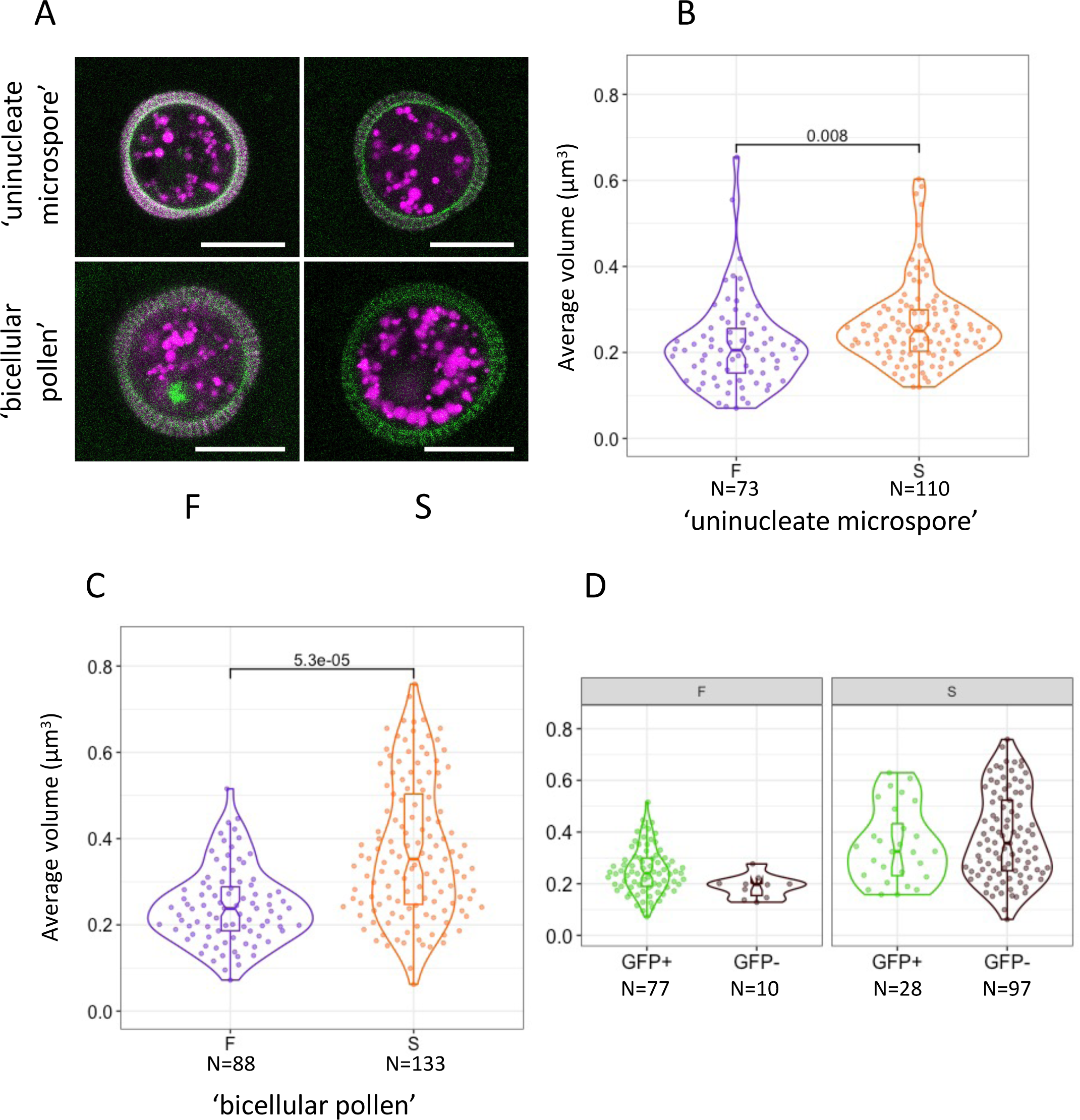
Mitochondria swelling in the pollen of sterile plants. A. Representative micrographs of pollen carrying the pBnUBX1::mtmCherry (magenta) and the pTip5;1::H2B-GFP (green) markers in the two analyzed stages of fertile (F) and sterile (S) plants. The images for the figure were produced by summing three slices of the original Z-stacks. Brightness and contrast were adjusted for visualization. Note that the pollen wall is visible due to autofluorescence. Scale bars: 10 µm. B, C. Average volume of mitochondria in pollen, according to the bud developmental stage (Table 2). B: ‘uninucleate microspore’ stage, C: ‘bicellular pollen’ stage. Each measurement point corresponds to an individual pollen grain. The pvalue of the likelihood ratio test comparing mixed linear models with or without considering the genotype as a variable is given above. D. Average volume of mitochondria in grains of the ‘bicellular pollen’ stage, according to the presence (GFP+) or not (GFP-) of the generative nucleus, based on the fluorescence of the germline marker. The complete dataset is listed in Table S8. F: [Kz-9]Cvi, S: [Sha]Cvi. N = number of measurement points.

## Discussion

Whereas *orf117Sha* did not present the known features often used to predict candidates for CMS genes (Kitazaki *et al*., 2023), it remained the best candidate after the assembly of Sha and Kz-9 mitochondrial genomic sequences. By producing and analyzing mitoTALEN-induced deletions in the Sha mitochondrial genome, we confirmed that *orf117Sha* is indeed the Sha-CMS causal gene. The mitochondrial deletions analyzed in mitoTALEN reverted plants resulted from recombination between copies of repeated sequences and were accompanied by the duplication of a large region of the mitochondrial genome which carried several conserved mitochondrial genes, including *cob*. These results are coherent with previously reported plant mitochondrial genomes modified by mitoTALEN (Kazama *et al*., 2019) and with the model of plant mitochondrial genome evolution proposed by Small and Leaver (1989).

The mitoTALEN-induced deletion approach is becoming the golden standard for the validation of CMS- candidate genes (Kazama *et al*., 2019; Omukai *et al*., 2021; Takatsuka *et al*., 2022; Kuwabara *et al*., 2022). Until recently, the production of male sterile plants by nuclear expression of the CMS-candidate gene fused to a mitochondrial-targeting sequence has been largely used to validate CMS-candidate genes. However, this approach is not fully reliable to validate a candidate CMS-gene, see for example the cases of T-CMS in maize (Chaumont *et al*., 1995) or Ogura-CMS in rapeseed (Duroc *et al*., 2006), where the CMS-inducing proteins produced from transgenes were incorrectly imported or located in the mitochondria. Still, provided that pollen death is robustly induced in transgenic plants by the same mechanism as in CMS ones, it could open opportunities for structure-function studies of CMS-inducing proteins. In the present study, only one out of 21 transgenic plants presented the expected pollen phenotype (half dead pollen), but this plant hardly transmitted the transgene to its selfing progeny, indicating a default in transmission through both the male and female germlines. As the *UBQ10* promoter was recently reported to be active in the embryo sac (Hu *et al*., 2023), this observation is compatible with a toxic effect of the transgene in both male and female gametophytic tissues. Noteworthily, this would imply that the toxicity of ORF117SHA is not efficient on sporophytic tissues, since transgenic plants presented no apparent phenotypical modifications. Indeed, a lack of transgene transmission was also reported for Arabidopsis plants with 50 % dead pollen, transformed with a mitochondrion-targeted ORF108 protein, which shares 56% identity with ORF117SHA (Gobron *et al*., 2013) and the authors concluded that ORF108 had a toxic activity in both male and female gametophytes (Kumar *et al*., 2012). In the present study, it is conceivable that the *UBQ10* promoter activity, which was not detectable before the bicellular pollen stage (Figure S1), was not sufficient to accumulate enough amount of ORF117SHA in the mitochondria of pollen at the proper stage to result in the phenocopy of the Sha-CMS. In this case, the use of a promoter that would be strongly active in the earliest stages of pollen development, yet to be identified, might be the key to obtaining a genuine phenocopy of the Sha-CMS and opening a path towards a structure-function study of the ORF117SHA protein.

Considering that CMS-induced pollen death is likely to rely on pollen-specific features, we focused our investigations on pollen-centered approaches, implementing recently developed imaging tools. Fluorescent microscopy of pollen expressing a nuclear-located GFP produced specifically in the male germline (Borg *et al*., 2011) allowed use to refine the developmental stage at which pollen death occurs in Sha-CMS plants. Our results clearly showed that the development of pollen grains is highly desynchronized in the sterile genotype. At least a part of the pollen grains was delayed in their development, as illustrated by the cooccurrence of grains with one or two GFP-marked nuclei at the ‘PMII’ stage. In addition, the stage of development ultimately reached before abortion was different amongst pollen grains. Therefore, oppositely to the HL-CMS of rice where the pollen cell cycle was reported to be arrested before PMII (Wang *et al*., 2013), in Sha-CMS pollen development does not stop at a precise stage but, depending on each pollen grain, between the uninucleate microspore and the trinucleate pollen stages. Grain-autonomy is the genetic rule in gametophytic CMS and it is consistent that it also applies for the death process, although this was not described previously, to the best of our knowledge.

We observed that as early as the ‘uninucleate microspore’ stage the average mitochondria volume of pollen was slightly higher in sterile compared to fertile plants (0.26 µm^3^ +/- 0.09 and 0.24 µm^3^ +/- 0.09, respectively). Due to the similarity between the compared genotypes, it is likely that this observation is linked to the presence of *orf117Sha*. Further investigations will be necessary to explore whether this observation is due to an impaired functioning of mitochondria in the sterile, which could possibly trigger the pollen death process. At the ‘bicellular pollen’ stage in sterile plants, the average volume of the pollen mitochondria became very heterogenous, with a bimodal distribution, a proportion of pollen grains showing mitochondria of average volume above 0.5 µm^3^. This spectacular morphologic change evoked the mitochondrial morphology transition observed in stress conditions such as mild heat shock or oxidative stress in protoplasts (Scott and Logan, 2008). The mitochondrial morphology transition was shown to precede cell death and proposed to be an indicator of the mitochondrial permeability transition (Scott and Logan, 2008). Mitochondrial permeability transition is an increased permeability of the inner mitochondrial membrane which leads to mitochondrial swelling, and has been described in plants, animals and microorganisms and associated with programmed cell death (Logan, 2008; Zancani *et al*., 2015; Ocampo-Hernández *et al*., 2022; Bernardi *et al*., 2023). It will be interesting to further investigate whether the mitochondrial permeability transition is involved in the swelling of mitochondria in the pollen of Sha-CMS plants. This will require the implementation of approaches for the monitoring of membrane potential in individual pollen mitochondria and the uptake into developing pollen of inhibitors/inducers of the mitochondrial permeability transition, such as lanthanum chloride or Ca^2+^. However, the coincidence between the heterogeneity of mitochondria volumes with the pollen death heterogenous timing leads us to hypothesize that mitochondria swelling likely reflects a late phase in the process of pollen abortion. No mitochondria swelling in pollen was reported in the gametophytic S-CMS of maize after mitochondria-targeted GFP imaging (Chamusco *et al*., 2022), and mitochondrial morphology was not described in the HL-CMS of rice, to our knowledge. However, changes in mitochondrial morphology were reported from electron microscopy observations in the tapetal cells of sporophytic sunflower PET1-CMS (Horner, 1977), rapeseed Ogura CMS (González-Melendi *et al*., 2008), soybean CMS (Smith *et al*., 2002), and of thermosensitive genic-male sterility of rice (Ku *et al*., 2003) and in pollen tubes experiencing the *Papaver rhoeas* self-incompatibility response (Geitmann *et al*., 2004), supporting the view that it is a convergent step in diverse cell death processes.

Although we did not identify the physiological trigger by which ORF117SHA induces the pollen death process, we were able to rule out that a deficit in ATP supply is the cause of pollen death by experimentally measuring Mg-ATP^2-^ content in pollen from fertile and sterile plants. However, our results do not rule out a role of the ATP synthase complex in the pollen death process. For example, ATP synthase dimers are thought to participate in the formation of the mitochondrial permeability transition pore (Zancani *et al*., 2020), provoking the mitochondrial swelling that precedes pollen death (Scott and Logan, 2008; Van Aken and Van Breusegem, 2015). Impairment of mitochondrial ATP synthase function has been linked to male sterility in several sporophytic CMSs, notably in the sunflower PET1-CMS (Sabar *et al*., 2003) and in maize CMS-C (Yang *et al*., 2022), where it involved the production of aberrant variants of ATP synthase subunits. Although these studies report lower ATP content in extracts from flower/anther tissues, we are not aware of a direct measure of ATP content in the tapetum. Another possible trigger to be tested remains the accumulation of ROS, which could induce an oxidative stress and lead to mitochondria swelling and pollen death, similarly to experimentally ROS-induced cell death described by Scott and Logan (2008). Indeed, ROS accumulation has been proposed to contribute in pollen death in the HL-CMS of rice (Wang *et al*., 2013), as well as in several sporophytic CMSs, such as cotton CMS (Jiang *et al*., 2007), pepper CMS (Deng *et al*., 2012), or peach CMS (Cai *et al*., 2021). Unfortunately, we could not address the accumulation of ROS in developing Arabidopsis pollen, due to the unfavorable ratio between the fluorescence of the ratiometric redox reporter roGFP2-GRX1 (Lukyanov and Belousov, 2014) and autofluorescence of the pollen wall. This emphasizes the need for efficient promoters to drive strong expression of biosensors in early developing pollen, for the imaging of pollen grains and proper analysis of the results.

The use of pollen-centered *in vivo* cytological approaches allowed us to unveil the grain-autonomous pollen abortion process in Sha-CMS plants: pollen development pace slows down independently in each pollen grain, probably as early as the uninucleate microspore stage; at one point, either before or after PMI, and rarely after PMII, the swelling of mitochondria would mark the no-return death process, either after the arrest of pollen development or causing it. Whether the early step of the process is functionally linked to the slight enhanced volume of the mitochondria in sterile at the uninucleate microspore stage remains to be investigated. Addressing the question of the triggering of pollen death by ORF117SHA will undoubtedly be facilitated by newly developed approaches for mitochondria purification from specific tissue/cell types, such as IMTACT (Boussardon *et al*., 2020). It will also require the enrichment of our toolbox with pollen-efficient promoters and with more markers or biosensors for the measure of key metabolites and the exploration of physiological parameters at the cell level. Furthermore, this will allow to address questions concerning the functional similarity between evolutionary related CMS-causing proteins such as ORF117SHA and ORF108.

## Supplementary data

The following supplementary data are available at JXB online.

*Table S1*. Illumina data statistics, mapping results and raw read accessions.

*Table S2*. Mitology pipeline parameters.

*Table S3*. Mitology pipeline results.

*Table S4*. PacBio data statistics, assembly results and contig accessions.

*Table S5*. Repeated sequences of at least 100 bp in Sha and Kz-9 mitochondrial genomes.

*Table S6*. Primers used.

*Table S7*. cATeam ratio data.

*Table S8*. Pollen mitochondrial volume data.

*Table S9*. Open reading frames ≥ 100 codons in Col-0, Kz-9 and Sha mitochondrial genomes.

*Table S10*. Analysis of mitoTALEN modified T1BC1 plants.

*Table S11*. Analysis of mitoTALEN modified T2BC1 plants.

*Fig. S1*. Expression profile of the *UBQ10* promoter during pollen development.

*Fig. S2*. Target sequences of mitoTalens in the *orf117Sha* coding sequence.

*Fig. S3*. Assessment of pollen developmental stages from pistil size.

*Fig. S4*. Expression profile of the pBnUBX1 promoter during pollen development.

*Fig. S5*. Annotated maps of *de novo* assembled Kz-9 and Sha chloroplast genomes. *Fig. S6*. Annotated maps of *de novo* assembled Kz-9 and Sha mitochondrial genomes. *Fig. S7*. PCR analyses of mitoTALEN modified T2BC1 plants

*Fig. S8*. IGV visualization of read coverage in the *cob* region in the sterile control [Sha]Cvi and the mitoTALEN revertants.

*Fig. S9*. Repair of mitoTALEN-induced DNA breaks through very short repeats.

## Acknowledgements

We thank Pr David Twell (Leicester University) for the sharing of the Tip5;1::GFP construct, Dr Sandrine Bonhomme and Dr Mathilde Grelon for their support and help during the preparation of the manuscript, Mrs Katia Belcram and Mrs Gladys Cloarec from the IJPB cytology platform for their valuable help in cytological approaches and confocal imaging, Mrs Nadia Bessoltane and Dr Fabienne Granier from the IJPB bioinformatic team for their support, Mrs Yoshiko Tamura for her efficient cloning of the mitoTALEN vectors and the internship students Mr Quentin Bodinier and Mr Victor Taieb for their help in analysis of transgenic plants and Mrs Myriam Shafie and Mrs Alexandrina Bodrug for their contribution to the development of the mitology pipeline. This work has benefited from the support of IJPB’s Plant Observatory technological platforms. We are grateful to the genotoul bioinformatics platform Toulouse Occitanie (Bioinfo Genotoul, https://doi.org/10.15454/1.5572369328961167E12) for providing computing and storage resources.

## Author contributions

ND, CB, MSc and FB: conceptualization; PA, DC, JT and FB: data curation; ND, CB, PA, DC, CGT, DB and FB: formal analysis; SiA, MSc and FB: funding acquisition; ND, CB, CGT, TN, AR, MSi and DV: investigation; PA, DC, TN and JT: methodology; FB: Project administration; AR, CC and SiA: resources; PA, DC, JT: software; CB, DC, TN, AR, MSi, JT, CC, SiA, MSc and FB: supervision; ND, CB, DC, MSc and FB: validation; ND, PA, DB, DV and FB: visualization; ND, PA, DC, DB and FB: writing original draft; ND, CB, PA, DC, DB, MSi, JT, CC, SiA, MSc and FB: writing review and editing.

## Data availability

The sequencing data described in this article are available at the European Nucleotide Archive (ENA) <http://www.ebi.ac.uk/ena/data/view> under the study PRJEB64929.

Bioinformatic codes developed in this work for sequence assemblies and validation are available at <https://github.com/jos4uke/mitology-pipeline.git>, <https://github.com/jos4uke/contigLink>, and <https://forgemia.inra.fr/ijpb-bioinfo/public/insilicorflp.git>.

The programs developed in this work for pollen and mitochondria segmentations are available at <https://doi.org/10.57745/56PALB>.

All other supplementary data are available at JXB online.

